# Dexamethasone-Induced p57-Mediated Quiescence Drives Chemotherapy Resistance in Sonic Hedgehog Medulloblastoma

**DOI:** 10.1101/2025.04.27.650711

**Authors:** Aryaman Sharma, Lauren Ellis, Kaytlin Andrews, Alexander Brown, Margaret T. Fuller, Matthew P. Scott, Teresa Purzner, James Purzner

**Author notes:** Corresponding Authors (JP), (TP).

## Abstract

Medulloblastoma (MB), the most common malignant pediatric brain tumor, remains difficult to cure upon relapse, with only 12.4% of patients surviving five years post-recurrence. While specific quiescent tumor cell populations are known to contribute to treatment-resistance, the molecular mechanisms that maintain quiescence remain poorly defined. Here, we identify the cell cycle inhibitor p57 as a regulator of quiescence and chemotherapy resistance in Sonic Hedgehog (SHH) MB. Nuclear p57 was enriched in Sox2+ and Nestin+ stem-like MB cells compared to proliferative Atoh1+ cells. Inducing p57 expression in SHH MB cells led to a six-fold increase in G_0_-phase cells and conferred resistance to the frontline chemotherapeutic vincristine. Clinically, dexamethasone is a glucocorticoid given to nearly all MB patients to manage cerebral edema and is administered with wide variability in timing and dosing. We show that dexamethasone significantly increased nuclear p57 levels and expanded the G_0_ population in both *Ptch1*^+/−^ and *Ptch1*^+/−^;*Trp53*^−/−^ SHH MB mouse models. Pre-treatment with dexamethasone reduced vincristine sensitivity in SHH MB cells. Together, our findings reveal a clinically relevant and previously unrecognized mechanism of treatment resistance, whereby dexamethasone, despite its benefits in managing edema, may inadvertently contribute to tumor persistence or recurrence by driving a quiescent, drug-resistant state. Addressing the lack of standardization in steroid use or targeting p57 may improve treatment response and reduce recurrence in patients diagnosed with SHH MB.

## INTRODUCTION

A major challenge in the treatment of medulloblastoma (MB), the most common malignant pediatric brain tumor, is disease recurrence. Approximately 30% of patients will experience tumor relapse, and despite aggressive salvage therapy, nearly all patients will succumb to recurrent disease (*1–3*). Developing targeted strategies to treat or prevent recurrent tumors requires a foundational understanding of the drivers of recurrence.

Medulloblastomas are classified into four subgroups: SHH, WNT, Group 3 and Group 4 (*4*). Though clonally distinct from their therapy-naïve primary tumors, recurrent tumors have been shown to retain their subgroup affiliation (*1, 5–8*). Each subgroup has a distinct recurrence pattern, suggesting underlying differences in etiologies. For example, SHH tumors exhibit a higher incidence of local recurrence relative to the more frequent metastatic recurrences observed with Group 3 and Group 4 tumors (*1, 8*). Temporally, SHH tumors, along with Group 3, recur more rapidly and are associated with shorter survival post recurrence compared to those in Group 4 (*1, 8*). Genetically, mutations in TP53, a tumor-suppressing transcription factor, are associated with highly aggressive tumors and occur exclusively in the SHH subgroup. (*3, 8, 9*).

Quiescent cancer stem cells are affiliated with recurrence in a wide range of tumors, including leukemia, melanoma, glioblastoma, and MB (*10–14*). Within SHH MB, a rare subpopulation of quiescent, multipotent progenitors, identified by accumulation of the transcription factor Sox2, has shown robust resistance to traditional therapies and the ability to cause tumor expansion upon transplantation (*14*). These quiescent, Sox2+ cells proliferate at a low frequency while retaining the capacity to differentiate into the Atoh1+ cells that comprise the majority of cells within the tumor (*14*). In contrast to their efficacy against proliferative cells, frontline chemotherapy, radiation and targeted SHH antagonists fail to kill quiescent cells, thereby enriching for the cell type most likely to cause recurrence (*14–16*).

The molecular machinery that maintains quiescence in Sox2+ cells remains poorly understood. However, the cyclin-dependent kinase inhibitor p57 is highly expressed in the rhombic lip, the region that gives rise to granule neuron precursors (GNPs), the developmental cell of origin of SHH MBs. p57 has also been identified as a key regulator of neural stem cell quiescence in the dentate gyrus of the adult hippocampus (*18, 19*).

Given the critical role of p57 in maintaining quiescent neural stem cells in the adult brain and its expression during GNP development, we tested whether p57 may also play a key role in maintaining quiescence in SHH MB. First, we investigate whether p57 levels vary in in Sox2+ cells relative to the tumor bulk. Next, we investigate whether elevated p57 expression is sufficient to drive SHH MB cells into a quiescent state and, as a results, incur greater resistance to frontline chemotherapy agents. Finally, we investigate whether dexamethasone, a corticosteroid commonly administered to manage tumor-associated edema, is capable of inducing p57-mediated quiescence and chemoresistance in SHH MB cells.

## RESULTS

### Sox2+ quiescent cells show enrichment of high nuclear p57 in SHH MB tumors

*Patched1* heterozygous (*Ptch1*^+/−^) murine models, which develop SHH MB tumors (Fig. 1A), were used to characterize p57 expression within SHH MBs. Low levels of p57 expression were observed throughout the tumor, with discrete subpopulations of cells exhibiting markedly higher protein levels (Fig. 1B). p57 protein colocalized with DAPI, consistent with its expected nuclear localization required for cell cycle inhibition (Fig. 1C). Nuclear p57 was significantly enriched in Sox2+ cells compared to Sox2-cells (Fig. 1D-E; p = 0.00028), in keeping with a potential role for p57 in regulating quiescence within SHH MBs.

**Figure 1:**
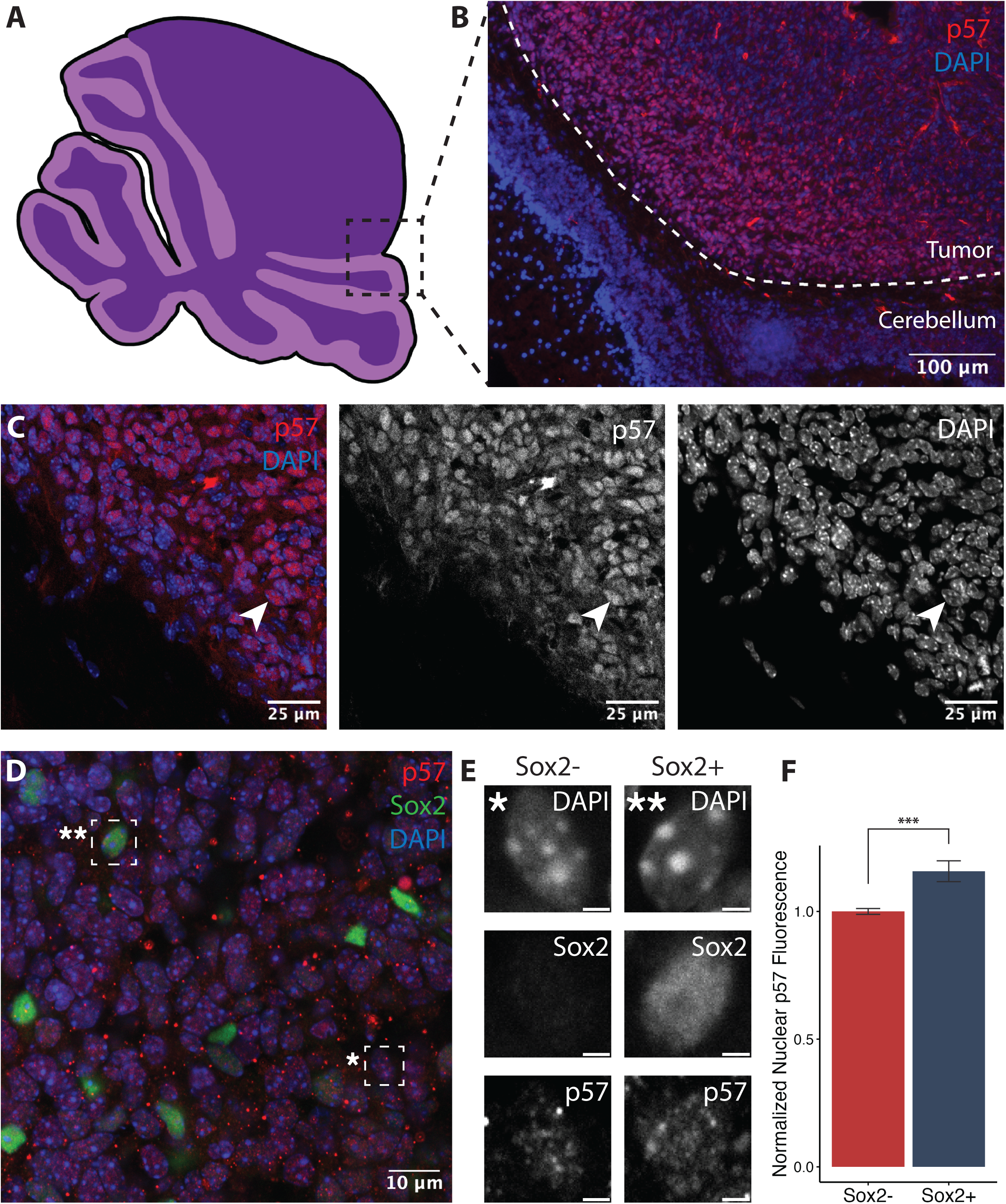
Subpopulations defined by high nuclear p57 in SHH medulloblastoma. **(A)** Schematic of a SHH medulloblastoma in *Ptch1*^+/−^ mice and magnified region of following confocal micrographs (dotted black box) **(B)** 10x and **(C)** 40x immunofluorescence image showing expression of nuclear p57 (red) and DAPI (blue) within MB **(D)** 60x high magnification immunofluorescence images showing colocalization of p57 (red) with Sox2 (green) in *Ptch1*^+/−^ MB **(E)** Single cell comparison of p57 immunofluorescence in Sox2-(‘) and Sox2+ (”) cells. Scale bars indicate 2µm **(F)** Quantification of p57 fluorescence showing significantly elevated p57 signal intensity in Sox2+ cells (n=667 nuclei) (***p < 0.001). P-values were calculated using Wilcoxon rank-sum test.

### p57 is enriched in Nestin+ and Sox2+ cells compared to Atoh1+ cells *in vitro*

When grown in culture, SHH MB cells from *Ptch1*^+/−^ mice (MB55) showed diffuse cytoplasmic p57 expression (Fig. 2). This was in contrast to IF images acquired from whole tumors that showed high nuclear p57 immunofluorescence (Fig. 1). Using high-throughput single-cell immunofluorescence imaging, p57 protein levels were quantified alongside Atoh1 (a marker of proliferative, transit-amplifying-like cells) and Sox2/Nestin (markers of quiescent, multipotent progenitors) (Fig. 2A-C). Consistent with its inhibitory role in cell cycle, we observed significantly higher levels of p57 protein in Nestin+ and Sox2+ cells relative to rapidly-cycling Atoh1+ cells (Fig. 2D; p < 0.005). p57 was predominantly localized in nuclei in Nestin+ cells, whereas Atoh1+ cells had a higher ratio of cytoplasmic to nuclear p57 (Fig. 2D; p = 0.032 and p = 0.037, respectively).

**Figure 2:**
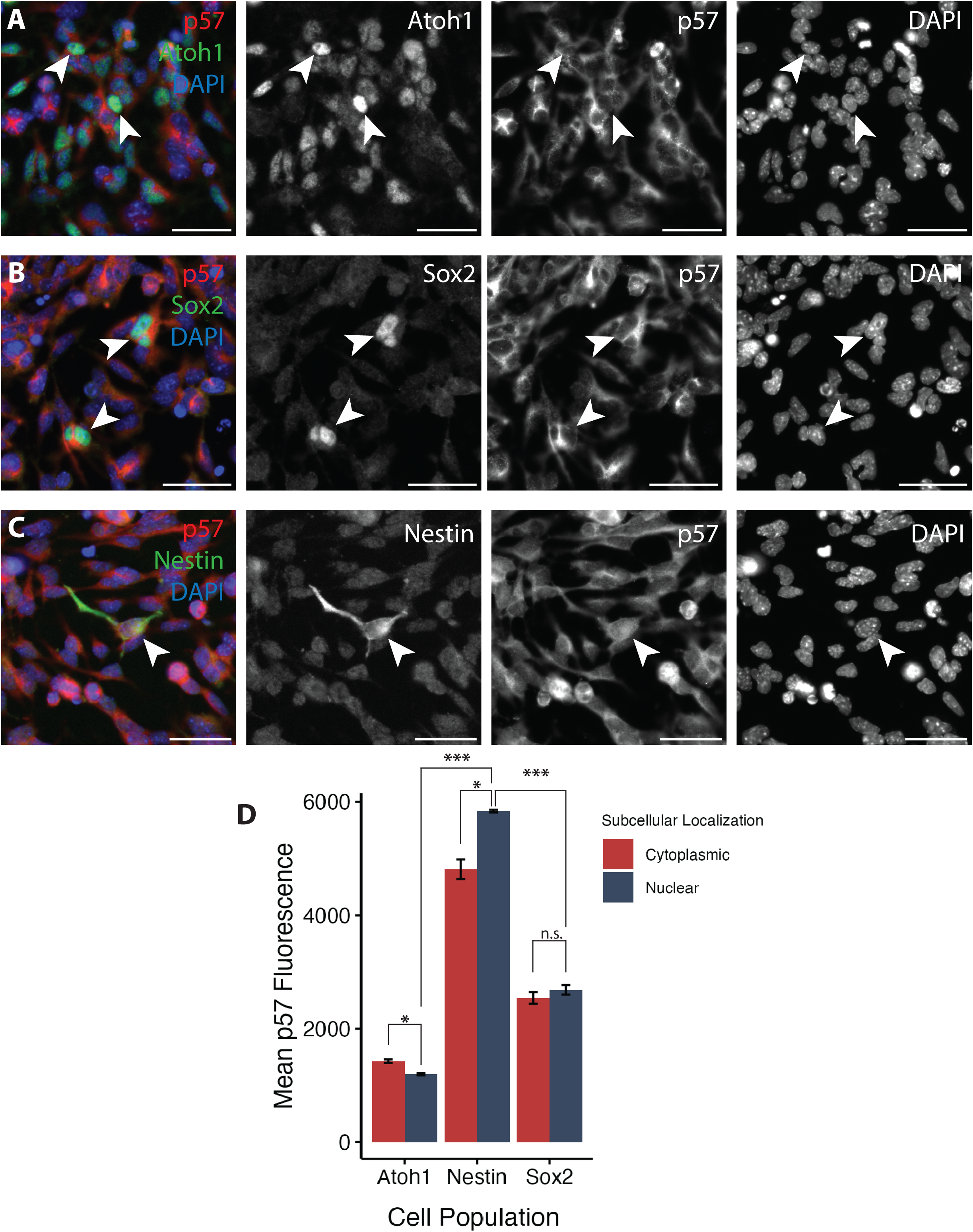
p57 is elevated in Sox2+ and Nestin+ stem-like SHH MB cells relative to fast cycling Atoh1+ MB cells. Representative images showing p57 colocalization with **(A)** Atoh1 (proliferative MB cell marker) **(B)** Sox2 and **(C)** Nestin (stem-like MB cell markers). Scale bars indicate 25µm **(D)** Quantification of p57 colocalization by marker and cytoplasmic (red bar) versus nuclear (blue bar) localization shows highest levels of p57 in Nestin+ cells followed by Sox2+ cells. Error bars represent SEM of triplicates. Across markers, p-values were calculated using one-way ANOVA followed by post-hoc Tukey’s HSD. Within markers, p-values of differences in cytoplasmic and nuclear fluorescence were assessed using paired t-test. n.s. = p > 0.05, *p < 0.05, **p < 0.001, ***p < 0.0001.

### Induced p57 leads to nuclear translocation and cell cycle arrest in SHH MB cells

Given the nuclear localization of p57 within Sox2+ cells, we next investigated whether p57 overexpression could induce increased nuclear localization of p57. An mCherry-DHFR-p57 fusion protein was virally transduced into a *Ptch1^+/−^* SHH MB cell line (*Ptch1*^+/−^;*E2fa>mCherry-DHFR-p57*) (Fig. 3A), resulting in constitutive production of the p57 protein fused to a destabilizing dihydrofolate reductase (DHFR) domain and a mCherry fluorescent protein. The DHFR domain drives rapid degradation of the fusion protein, resulting in moderate levels of p57 despite continuous transcription of the gene. Adding trimethoprim (TMP) stabilizes the DHFR domain in a rapid, reversible and dose-dependent manner, allowing a controllable increase in total p57 protein abundance (Fig. 3A). Treatment with increasing doses of TMP resulted in a significant increase in nuclear p57, as detected by single-cell imaging (Fig 3B-C). Cells treated with 5µM TMP to induce high p57 protein levels (Fig 3E, blue bars) were enriched in the G_0_ phase relative to uninduced controls (construct, but no TMP) (Fig 3E, red bars; 28.89% vs 11.75%, p = 0.144). High p57 cells, defined as TMP-treated cells with p57 levels exceeding those observed in control cells, had a 6-fold enrichment of cells in G_0_ (Fig. 3E – purple bars; 59.73% vs. 11.75%, p = 0.0012). Our data suggest that the high levels of p57 appear to be sufficient to induce cell cycle arrest in SHH MB tumor cells.

**Figure 3:**
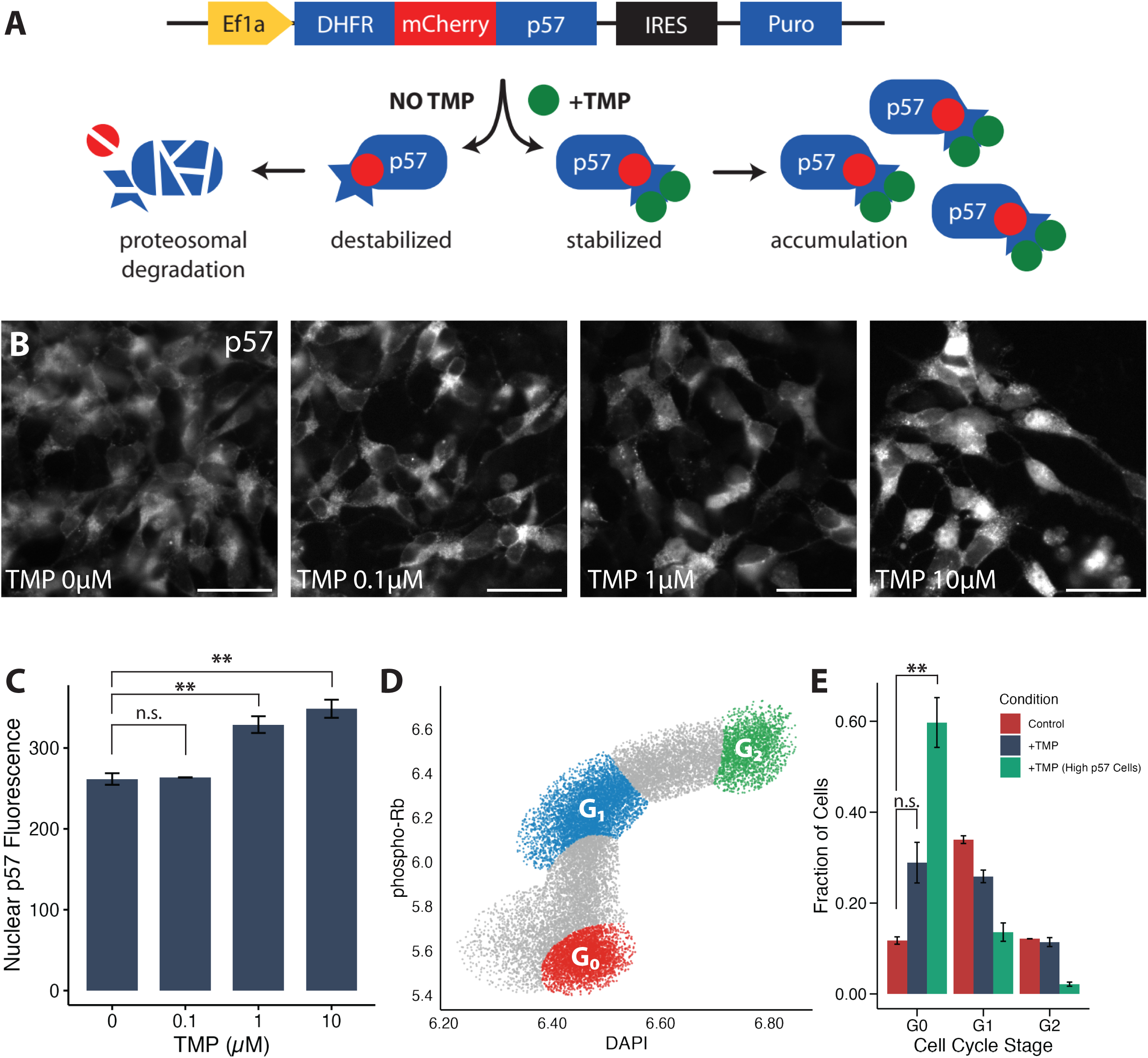
Inducing p57 protein abundance is sufficient to cause nuclear localization and cell cycle arrest in SHH MB cells. **(A)** Schematic representation of the DHFR-mCherry-p57 degron construct used to induce p57 protein **(B)** Representative images of mCherry-p57 expression responding to increasing doses of TMP. Scale bars indicate 25µm **(C)** Dose-dependent increase of nuclear p57 fluorescence in response to TMP escalation **(D)** Cell cycle clusters as measured by nuclear content (DAPI) and phospho-Rb fluorescence demarcating G_0_ (red), G_1_ (blue) and G_2_ (green) stages **(E)** Fraction of cells per cell cycle stage in control (red), TMP-treated (blue) and TMP-treated, high p57 (green) isolated cells. Error bars represent SEM across n=3 replicates (control) or n=4 replicates (TMP and TMP, High p57). P-values calculated using one-way ANOVA followed by post-hoc Tukey’s HSD. n.s. = p > 0.05, **p < 0.001.

### Elevated p57 confers resistance to vincristine in SHH MB cells

To determine whether high levels of p57 protect against frontline therapies, we used high content imaging to assess the viability of p57-DHFR SHH MB cells (*Ptch1*^⁺/⁻^;*Ef1a*>DHFR-mCherry-p57) treated with vincristine. Cells were either induced to express p57 (construct + TMP) or left uninduced as controls (construct without TMP). As expected, uninduced p57-DHFR cells demonstrated a dose-dependent decrease in cell viability when treated with vincristine, with 100% cell death at a 10uM dose (Fig. 4A). In contrast, while p57-DHFR cells treated with TMP (5uM) also demonstrated a dose-dependent decrease in cell viability when treated with vincristine, 22.4% of cells survived at vincristine doses as high as 10µM (Fig. 4A-B). Importantly, this treatment-resistant cell population was enriched for cells with high-p57 expression (Fig. 4B) and G_0_-phase cells (Fig. 4C).

**Figure 4:**
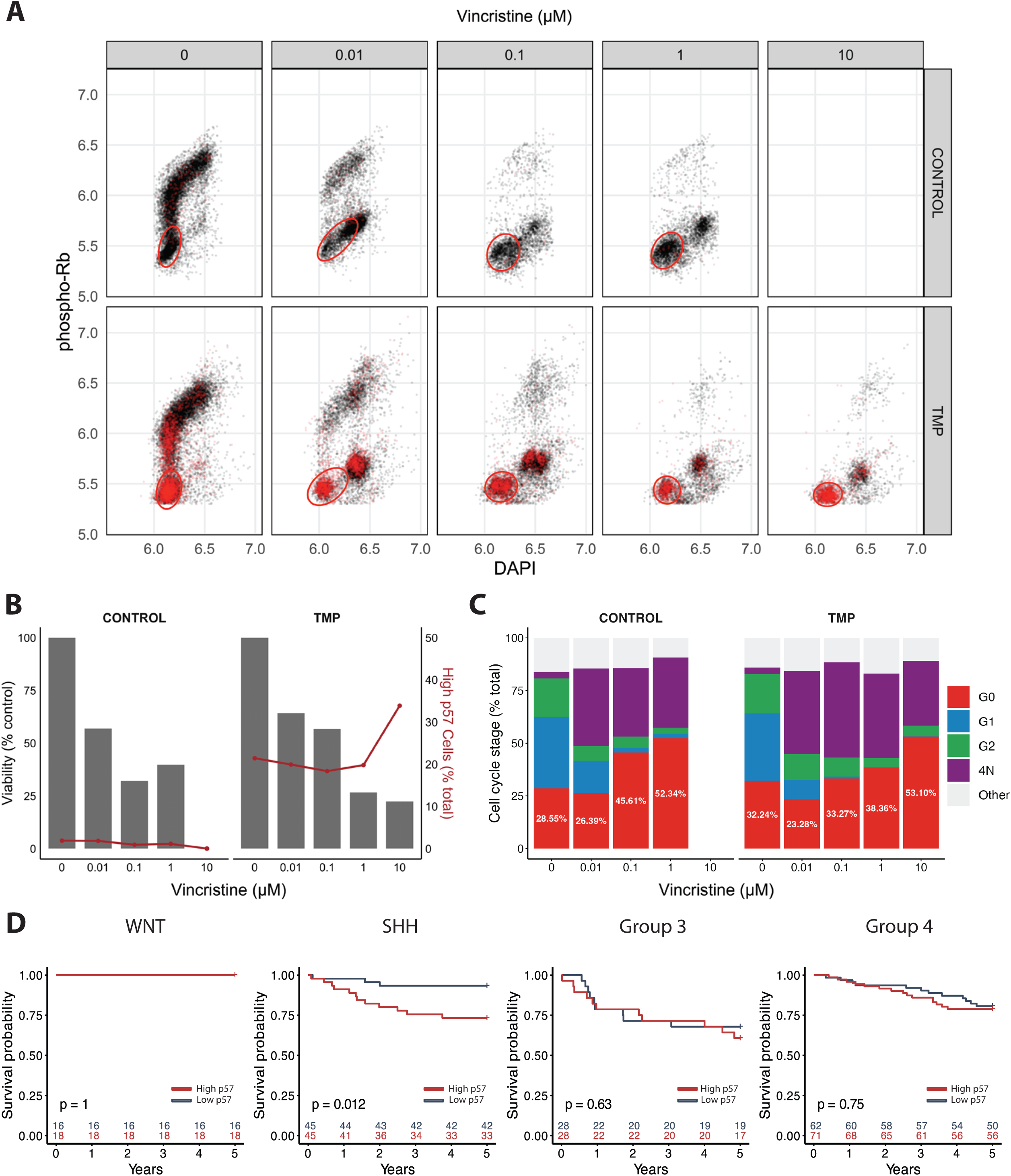
Elevated p57 confers protection from vincristine and correlates with poor survival in SHH MB. **(A)** DAPI-pRb plots showing control cells versus cells treated with 5µM TMP to induce high p57 in response to increasing doses of vincristine. Cells with high nuclear p57 are shown in red. The large red circle encloses G_0_ arrested cells **(B)** Viability (% control) (left axis) and High-p57 cells (% total) (right axis) in response to vincristine in control versus p57-induced cells. **(C)** Quantification of cell cycle phases shows failure to eliminate G_0_-phase cells across both induced and control groups **(D)** Kaplan-Meier curves comparing 5-year survival between high and low CDKN1C expression across MB subgroups.

To assess clinical relevance of p57 in medulloblastoma, we analyzed survival of patients stratified by p57 transcript (*CDKN1C*) levels. Elevated *CDKN1C* expression was associated with significantly lower 5-year survival rates for patients diagnosed with SHH-subtype medulloblastomas (93.3% vs 73.3%, p=0.012), but not for WNT (p=1), Group 3 (p=0.63) or Group 4 tumors (p=0.75) (Fig. 4D).

### Dexamethasone induces p57-mediated quiescence and chemoresistance in SHH MB cells

Given the potential role of p57 in mediating quiescence and chemoresistance, we next explored clinically-relevant regulators of p57 in SHH MB cells. Dexamethasone is a potent corticosteroid widely used for the management of brain tumor-associated edema. However, dexamethasone is also a known inducer of p57 expression via a glucocorticoid response element upstream of the *CDKN1C* gene (*20, 21*).

To examine whether dexamethasone treatment induces p57-mediated quiescence in SHH MB cells, cultured *Ptch1*^+/−^ SHH MB cells (IPM131) were treated with increasing doses of dexamethasone or vehicle control (0.01% DMSO) for 24 hours. Cells treated with 100nM dexamethasone had a 37% increase in p57 protein levels and an 8% increase in the fraction of cells in G_0_ relative to controls, as measured by high content imaging (Fig. 5A-C; p = 0.003 and p = 0.002). This effect was even more pronounced in *Ptch1^+/−^*;*Trp53^−/−^*SHH MB cells (PTCP53 304), where dexamethasone induced a 38% increase in p57, and a doubling of the fraction of cells in G_0_ after 24 hours (Fig. 5A-C; p = 0.009 and p < 0.001). These findings are particularly significant given that SHH MB tumors harboring *TP53* mutations are classified as very-high-risk and associated with poor prognosis. Finally, SHH MB cells pre-treated with 1µM dexamethasone for 24 hours exhibited a trend towards increased resistance to vincristine relative to controls, suggesting that dexamethasone may confer a protective effect against chemotherapy in these cells (Fig. 5D). These results demonstrate that dexamethasone-induced p57 expression promotes quiescence in SHH MB cells, thereby protecting them against frontline therapies including chemotherapy and radiation.

**Figure 5:**
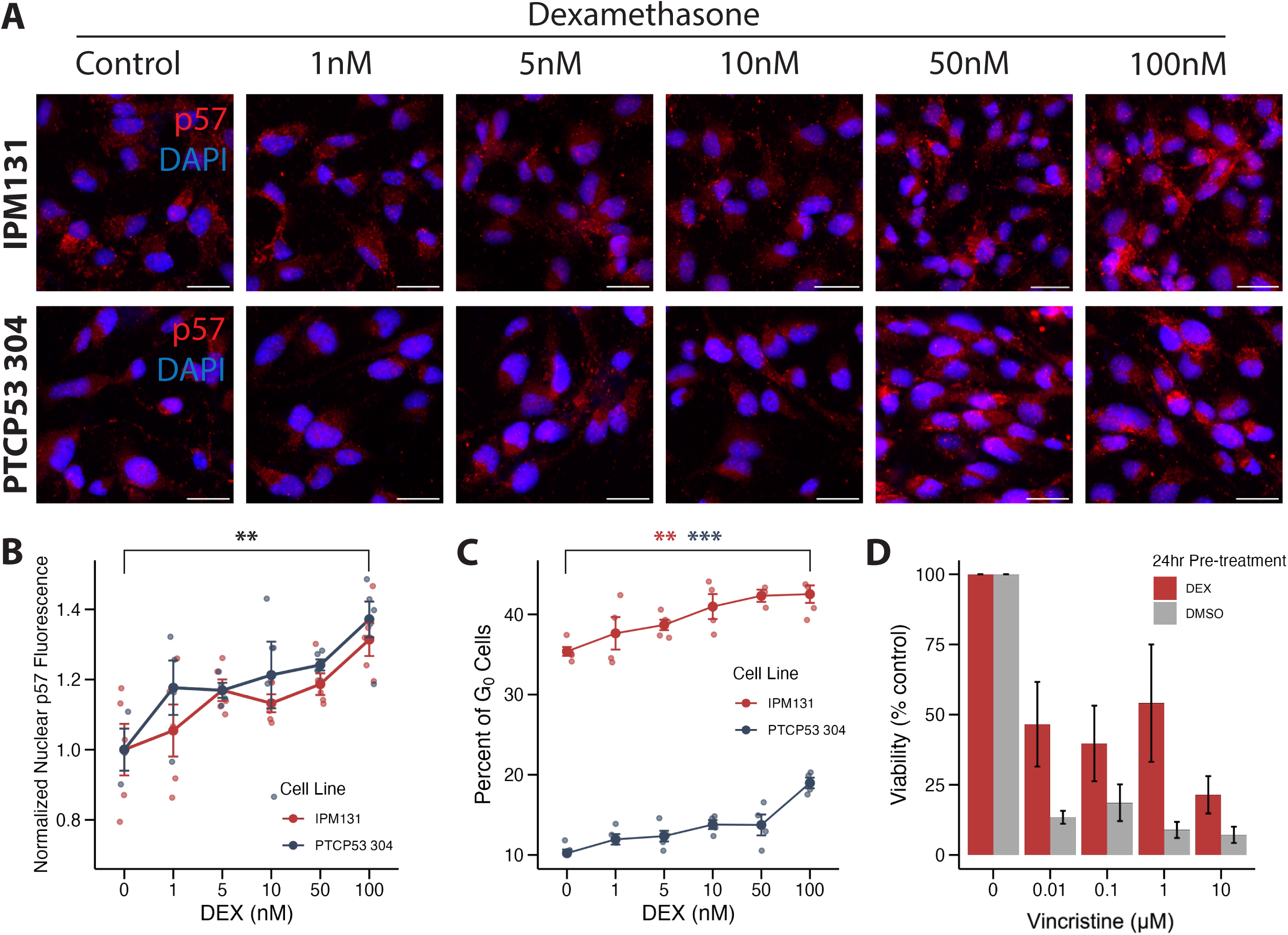
Dexamethasone induces p57-mediated cell cycle arrest and resistance to vincristine. **(A)** Representative immunofluorescence images showing p57 (red) and DAPI (blue) staining in *Ptch1^+/−^* (IPM131) and *Ptch1^+/−^;Trp53^−/−^* (PTCP53 304) SHH MB cells treated with increasing doses of dexamethasone. Scale bars represent 25µm **(B)** Quantification of nuclear p57 fluorescence intensity and **(C)** percentage of G_0_ cells relative to control following dexamethasone treatment **(D)** Quantification of cell viability in control and SHH MB cells pre-treated with 1 µM dexamethasone for 24 hours followed by vincristine exposure. Error bars represent SEM across five replicates. P-values calculated using one-way ANOVA followed by post hoc Tukey’s HSD. n.s. = p > 0.05, *p < 0.05, **p < 0.001, ***p < 0.0001

## DISCUSSION

Here we report a novel mechanism of quiescence and chemoresistance in SHH medulloblastoma. We first characterized p57 expression in SHH MB and found that p57 is relatively enriched in known quiescent MB cell populations, both *in vivo* and *in vitro*. We then show that over-expression of p57 is sufficient to induce G0 arrest and chemoresistance. In keeping, chemoresistant cells are enriched for high p57 expression levels and G0 arrest. Together, these findings suggest that p57 may support the survival of a subpopulation of SHH MB cells that evades current frontline therapies which target proliferative cells. The persistence of quiescent cancer cells during therapy may enable them to re-enter the cell cycle at a later time, seeding recurrence and undermining long-term treatment efficacy. These experimental findings are supported by clinical observations: high *CDKN1C* expression is associated with poorer survival among patients in the SHH subgroup, but not other MB subgroups. Finally, we demonstrate that dexamethasone increases p57 levels and induces quiescence in SHH MB cells, a finding with immediate clinical implications. Dexamethasone is a potent corticosteroid widely used for the management of brain-tumor related edema, typically administered both before and during chemotherapy and radiation treatments (*22*). However, a lack of systematic evidence on optimal dosing regimens has led to significant variability in its administration, often guided more by clinician preference than evidence-based practices (*23, 24*). Our findings provide a compelling reason for the urgent need for standardized and judicious use of dexamethasone in patients receiving adjuvant therapy. The dual effects of dexamethasone – increased p57 and tumor cell quiescence – were particularly pronounced in Trp53-mutant cells. SHH MB tumors carrying TP53 mutations are associated with treatment resistance and, as a result, particularly poor prognosis. Our data suggest that dexamethasone may inadvertently contribute to treatment resistance by supporting a population of quiescent, therapy-resistant cells. These results highlight the need to carefully assess the use of dexamethasone in patients diagnosed with SHH MB, especially in high-risk tumors, and to optimizing its dosing to avoid undermining the efficacy of frontline therapies.

Collectively, our findings identify p57 as a novel mechanism of quiescence in SHH MB and as a mediator of therapy resistance. This study not only provides insight into the role of p57 in inducing quiescence, but also raises important questions about how quiescent cells contribute to treatment resistance and subsequent tumor recurrence. A deeper understanding of how quiescence is regulated and maintained is critical for the development of novel therapies that can effectively target quiescent tumor cells and improve patient outcomes for this devastating disease.

## MATERIALS AND METHODS

### Mice

Mice heterozygous for *Patched1* (*Ptch1*^+/−^) spontaneously form SHH medulloblastomas(*25, 26*). Mice with developing MBs were identified from their neurological decline. Animals were monitored for ataxia, hunching and immobility. When neurologic decline was noted, the animals were euthanized and brains bearing SHH MBs were prepared for cryosectioning. All animal studies were approved by the Stanford APLAC review board protocol.

### Medulloblastoma cell culture

Murine SHH MB lines were grown either in suspension as neurospheres or adhered to coated plates. *Ptch1*^+/−^ MB cell lines MB21, MB55, and MB56 were generated in the Segal lab (Dana Farber Cancer Institute, Harvard Medical School) and grown as neurospheres while IPM131 and the *Ptch1*^+/−^;*Trp53*^−/−^ MB cell line, PTCP53 304, generated in the Dirks lab (Brain Tumour Research Centre, The Hospital for Sick Children) were grown adherently. Neurospheres (MB21, 55 and 56) were maintained in DMEM/F12 (Gibco, A4192001) supplemented with B-27 minus vitamin A (Gibco, 12587010) and 1% penicillin-streptomycin (Gibco, 15070063). Adherent cultures were maintained in Neurocult Basal Medium (Mouse & Rat) (Stemcell Technologies, 05700) supplemented with B-27 minus vitamin A, N-2 (Gibco, 17502048), 2mM L-glutamine (Sigma, G7513), 75µg/ml BSA (Sigma, A7030), 10µg/ml rhEGF (Stemcell Technologies, 78006), 10µg/ml rhFGF (Stemcell Technologies, 78003), 2µg/ml Heparin (Stemcell Technologies 7980). Adherent cultures were grown on Corning Costar 6-well cell culture plates (Corning, 3516) by coating them with 0.01% poly-L-ornithine (PLO) (Sigma, P4957) for 30 minutes at room temperature followed by 5µg/ml laminin (Sigma, L2020) in PBS overnight at 37°C.

### High content immunofluorescence imaging

Cell culture imaging for MB cells was performed in 96-well, glass bottom plates (Cellvis P96– 1.5H-N). Plates were coated with PLO for 30 minutes at room temperature followed by 10µg/ml laminin in PBS overnight at 37°C. Cells were passaged and dissociated using Accutase (Sigma, A6964) and filtered through a 70µm cell strainer to remove large aggregates before seeding at a density of 10,000 cells per well. Cells were allowed to adhere overnight before treatment with trimethoprim, vincristine, dexamethasone, or their combinations the following day. Following treatment, cells were fixed with 4% paraformaldehyde for 15 minutes at room temperature.

Adhesion to non-target proteins by antibodies was blocked by incubating cells for one hour with 5% serum, 1% BSA and 0.2% Triton X-100 for 1 hour at room temperature. Primary antibodies were then applied, and the preparations were incubated overnight at 4°C. Primary antibodies used in this study are described in Table 1. Donkey anti-immunoglobulin G (IgG) secondary antibodies against mouse, goat or rabbit conjugated to Alexa-488, Alexa-594 and Alexa-647, respectively, were prepared at 1:500 (Jackson Immunoresearch) and incubated for 1 hour at room temperature. High-throughput image acquisition was performed on the ImageXpress Micro XLS Widefield High Content Screening System (Molecular Devices, Sunnyvale, CA) or the Mica Microhub (Leica Microsystems, Wetzlar, Germany) using 20x Nikon objectives. High-throughput single-cell fluorescence data was generated as previously described using custom MATLAB scripts (*27*). Briefly, images of nuclei were segmented using DAPI, followed by extraction of fluorescence intensity values for each nucleus. For cytoplasmic quantification,

**Table 1:** Primary antibodies.

| Antibody | Species | Supplier | Catalog | Dilution |
| --- | --- | --- | --- | --- |
| p57 | Rabbit monoclonal | Abcam | ab4058 | 1:200 |
| p57 | Rabbit polyclonal | Sigma | SAB4500071 | 1:200 |
| Sox2 | Rabbit polyclonal | Abcam | ab97959 | 1:500 |
| Sox2 | Goat polyclonal | R&D Systems | AF2018 | 1:300 |
| phospho-Rb<br>(Ser<sup>807/811</sup>) | Rabbit monoclonal | Cell Signaling<br>Technology | 8516 | 1:1000 |
| Nestin | Mouse monoclonal | Invitrogen | MA1-110 | 1:100 |
| Atoh1/Math1 | Mouse monoclonal | Developmental<br>Studies<br>Hybridoma Bank |  | 1:50 |

Voronoi cells – geometrically defined regions around each nuclear centroid based on proximity – were generated to approximate individual cell boundaries. Cytoplasmic channel signals were then measured using adaptive thresholding, generating a cytoplasmic mask from which pixel intensity data was determined. Downstream analysis and figure generation was performed in R.

### Cell cycle classification and marker identification

Following segmentation and single-cell immunofluorescence quantification, cell cycle analysis was performed in R using the mclust package. Cells were clustered with Gaussian finite mixture modeling using log_10_-transformed median intensity of DAPI, phosphorylated Rb and cell area to demarcate G_0_, G_1_, and G_2_ cell states. For *in vitro* cell classification of Nestin+, Sox2+ and Atoh1+ cells, nuclear intensity (DAPI) of each cell was plotted along a histogram and manual thresholds were set at the saddle points of the resulting bimodal distributions. Comparative and subcellular localization analysis was conducted on marker-positive cells.

### Immunohistochemistry and Sox2+ cell identification

Brains from mice with MBs were perfused with 4% PFA and fixed for 16 hours. Brains were washed with PBS overnight and placed in a sucrose gradient (15% and 30%) until the tissue sank. Fixed whole brains were embedded in O.C.T., cryosectioned at 15µm, and stored at −20°C until ready to be stained. For high magnification staining, sections underwent heat-induced epitope retrieval by heating slides in 10mM citrate buffer at pH 6 and heating at 95°C for 20 minutes. Background binding sites in the sections were then blocked by incubation with serum, followed by primaries and secondaries, and counterstaining as described above. High magnification images were acquired on a Nikon AX confocal microscope (Nikon Instruments Inc., Melville, NY) using a 60x oil immersion objective. High-throughput single-cell image quantification was performed using a custom MATLAB similar to our *in vitro* single-cell imaging script **(High content immunofluorescence imaging)**, whereby DAPI was used to segment nuclei followed by the extraction of pixel intensity values for each nucleus. A manual threshold at the saddle point of the bimodal Sox2 intensity distribution to distinguish Sox2+ from Sox2-cells, allowing comparison of pixel intensities in both populations.

### DNA constructs

To generate the mCherry-DHFR-p57 fusion protein gene, cDNA was synthesized from RNA extracted from cerebellar granule neuron precursors isolated from a C57BL/6 background. Reverse transcription was performed using the Superscript III First-Strand Synthesis Supermix (Thermo Fisher, 18080400). The *Cdkn1c* (p57) insert was PCR-amplified from cDNA using Q5 High-Fidelity DNA Polymerase (New England Biolabs, M0491) with primers designed to introduce overlapping sequences compatible with DHFR-mCherry sequences. The DHFR-mCherry fragment was amplified from EF1a-DHRF-mCherry-ARHGAP29, a gift from Dr. Arnold Hayer, using primers designed for subsequent Gibson Assembly. PCR products were gel-purified using a MinElute Gel Extraction Kit (Qiagen, 28604). The pLV-EF1a-IRES-Puro plasmid backbone was digested with BamHI-HF and EcoRI-HF (NEB, RO136 and R3101), followed by dephosphorylation with Calf Intestinal Phosphatase (NEB, M0525) for 30 minutes. For assembly, Gibson Assembly, 2X Master Mix (NEB, M0270) was used to insert the DHFR-mCherry and p57 fragments into the linearized pLV-EF1a-IRES-Puro backbone. Following the Gibson reaction, competent NEB 5-alpha E. coli cells were transformed.

### Lentivirus

Lentivirus was produced in HEK293T cells using a second-generation packaging system. 293T cells were plated at 2 × 10^6^ cells per 10 cm dish one day before transfection in DMEM supplemented with 10% fetal bovine serum (FBS) but without Penicillin-Streptomycin. Lipofectamine 2000 (Thermo Fisher, 11668027) was used for transfection. A total of 22 µg of plasmid DNA was transfected per plate, consisting of 10 µg of pLV-DHFR-mCherry-p57-IRES-Puro, 4 µg of VSVG envelope plasmid, and 8 µg of psPAX2 (packaging plasmid, Δ8.92). DNA was diluted in Opti-MEM (Thermo Fisher, 31985062) with P3000 reagent (2 µL per µg DNA) (Thermo Fisher, L3000001) and incubated with Lipofectamine 2000 for 15 minutes before being added dropwise to the cells. After 6 hours, the transfection media was replaced with fresh DMEM supplemented with 10% FBS and 10 mM HEPES to stabilize the viral particles. Viral supernatant was collected at 48 hours and stored at 4°C, followed by a second collection at 96 hours post-transfection. The combined supernatants were centrifuged at 1000 rpm for 5 minutes to remove cellular debris, followed by filtration through a 0.45 µm low-protein binding filter. Lentiviral particles were concentrated using Lenti-X Concentrator (Takara Bio, 631231) overnight at 4°C and resuspended in PBS. For transduction, *Ptch1^+/−^* SHH MB55 cells were plated in 6-well plates at 2 × 10⁵ cells per well one day before infection. Lentiviral concentrate was added to the media with 2 µg/mL polybrene (Sigma, TR-1003) to enhance infection efficiency. After overnight incubation, the media was exchanged with fresh, virus- and polybrene-free media. For selection, cells were cultured in 0.5 µg/mL puromycin starting 48 hours post-transduction. Non-transduced control cells were completely eliminated after 5 days, while p57-expressing cells survived and formed spheres.

### Human Kaplan-Meier curves

Human Kaplan-Meier curves were generated using the Cavalli cohort of 763 primary human medulloblastoma samples (*28*). Samples were first stratified by MB subgroup affiliation. Within each subgroup, samples with high and low p57 expression were defined as the top and bottom 25% of CDKN1C-expressing samples, respectively. Survival curves were estimated using the Kaplan-Meier method using the survfit package in R.

## Author contributions

Conceptualization: AS, AB, MTF, MPS, TP and JP

Methodology: AS, AB, MTF, MPS, TP and JP

Investigation: AS, AB, LE, KA, TP and JP

Funding acquisition: JP and TP

Supervision: JP, TP

Writing – original draft: AS, KA, AB, MTF, MPS, TP and JP

## Data and materials availability

Custom analysis scripts are available following Github repositories; jpurzner/cell_culture_segment

