## Supplemental Figures for "Dexamethasone-Induced p57-Mediated Quiescence Drives Chemotherapy Resistance in Sonic Hedgehog Medulloblastoma"

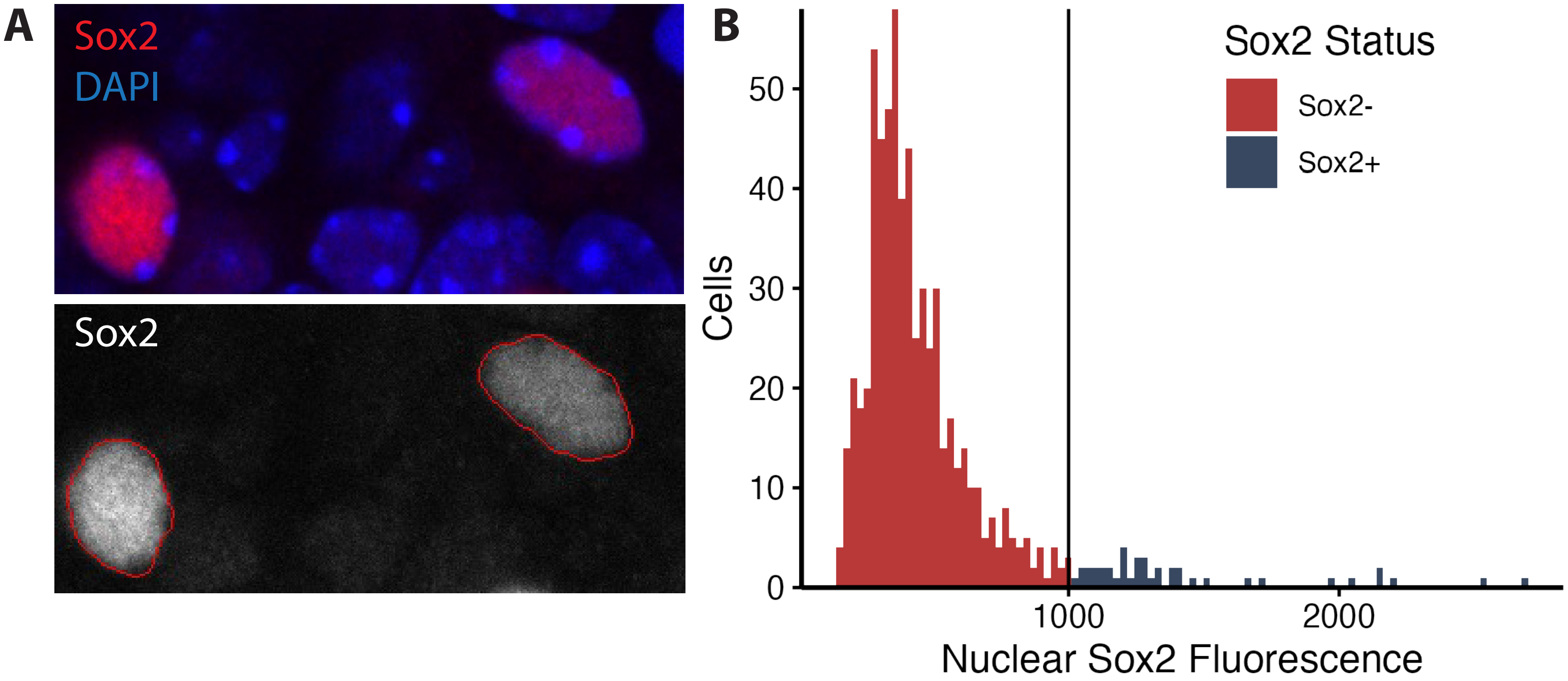
A
Sox2
DAPI
Sox2
B
Cells
50
40
30
20
10
0
1000
2000
Nuclear Sox2 Fluorescence
Sox2 Status
Sox2-
Sox2+

**Supplementary Figure 1: Validation of automatic identification of Sox2+ cells in *Ptch1*$^{+/-}$ SHH MB.** (A) Representative 60x immunofluorescence image of *Ptch1*$^{+/-}$ MB showing Sox2 (red) and DAPI (blue) (B) Representative detection of Sox2+ nuclei (C) Threshold used to delineate Sox2- (red) from Sox2+ (blue) nuclei

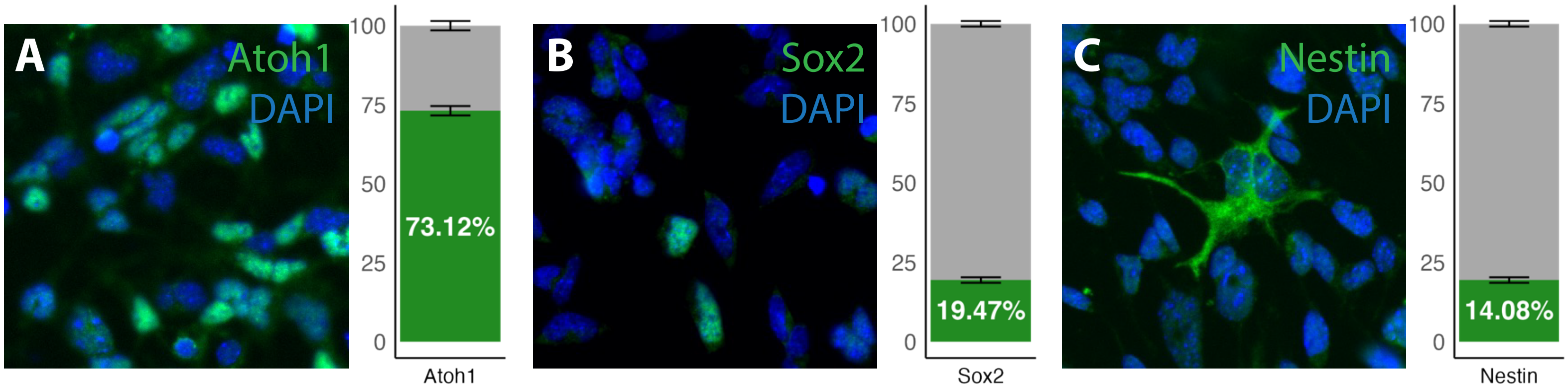
A
Atoh1
DAPI
100
75
50
25
0
73.12%
Atoh1
B
Sox2
DAPI
100
75
50
25
0
19.47%
Sox2
C
Nestin
DAPI
100
75
50
25
0
14.08%
Nestin

**Supplementary Figure 2: Relative proportions of MB cell populations in culture.**

Representative 20x micrographs and relative proportions of (A) Atoh1+, (B) Sox2+, and (C) Nestin+ *Ptch1*$^{+/-}$ MB cells in culture

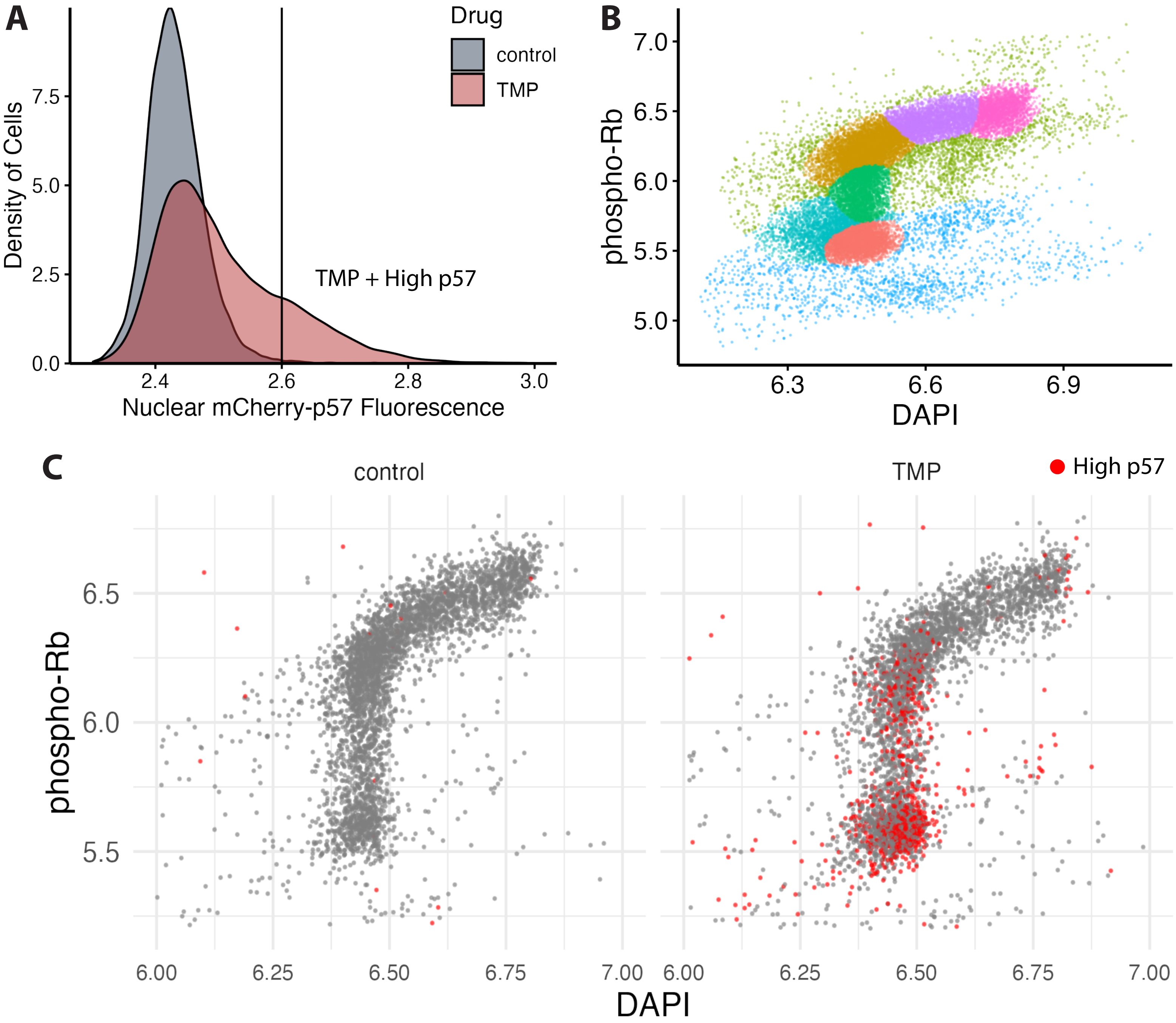
A
Drug
control
TMP
Density of Cells
7.5
5.0
2.5
0.0
2.4
2.6
2.8
3.0
Nuclear mCherry-p57 Fluorescence
TMP + High p57
B
phospho-Rb
7.0
6.5
6.0
5.5
5.0
6.3
6.6
6.9
DAPI
C
control
TMP
High p57
phospho-Rb
6.5
6.0
5.5
6.00
6.25
6.50
6.75
7.00
DAPI

**Supplementary Figure 3: Validation of TMP inducible p57 degron construct in Ptch1+/- MB cells** (A) Nuclear mCherry-p57 signal of control versus TMP-treated cells. The vertical line demarcates TMP-treated cells with higher p57 expression than the maximum observed in control cells (B) Raw clusters from density-based modeling (C) Enrichment of high p57 cells (red) in the $G_0$ cluster as detected by immunofluorescence.

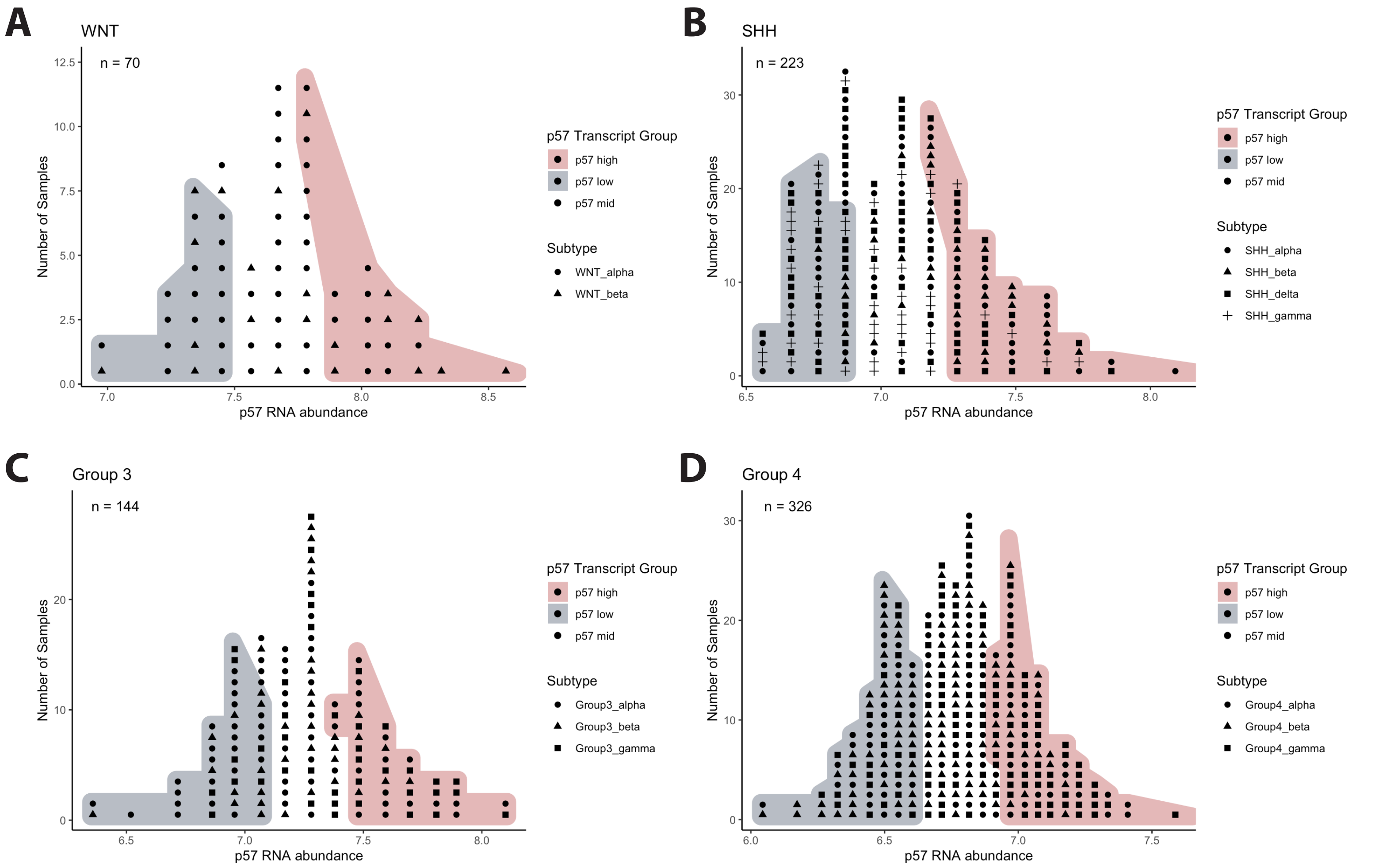
A
WNT
n = 70
Number of Samples
p57 RNA abundance
0.0
2.5
5.0
7.5
10.0
12.5
7.0
7.5
8.0
8.5
p57 Transcript Group
p57 high
p57 low
p57 mid
Subtype
WNT_alpha
WNT_beta
B
SHH
n = 223
Number of Samples
p57 RNA abundance
0
10
20
30
6.5
7.0
7.5
8.0
p57 Transcript Group
p57 high
p57 low
p57 mid
Subtype
SHH_alpha
SHH_beta
SHH_delta
SHH_gamma
C
Group 3
n = 144
Number of Samples
p57 RNA abundance
0
10
20
6.5
7.0
7.5
8.0
p57 Transcript Group
p57 high
p57 low
p57 mid
Subtype
Group3_alpha
Group3_beta
Group3_gamma
D
Group 4
n = 326
Number of Samples
p57 RNA abundance
0
10
20
30
6.0
6.5
7.0
7.5
p57 Transcript Group
p57 high
p57 low
p57 mid
Subtype
Group4_alpha
Group4_beta
Group4_gamma

**<u>Supplementary Figure 4: Histogram of p57 RNA abundance</u>** (Cavalli, 2017) demarcating high (red) and low p57 (blue) expressing samples for (A) WNT, (B) SHH, (C) Group 3 and (D) Group 4 subtypes.

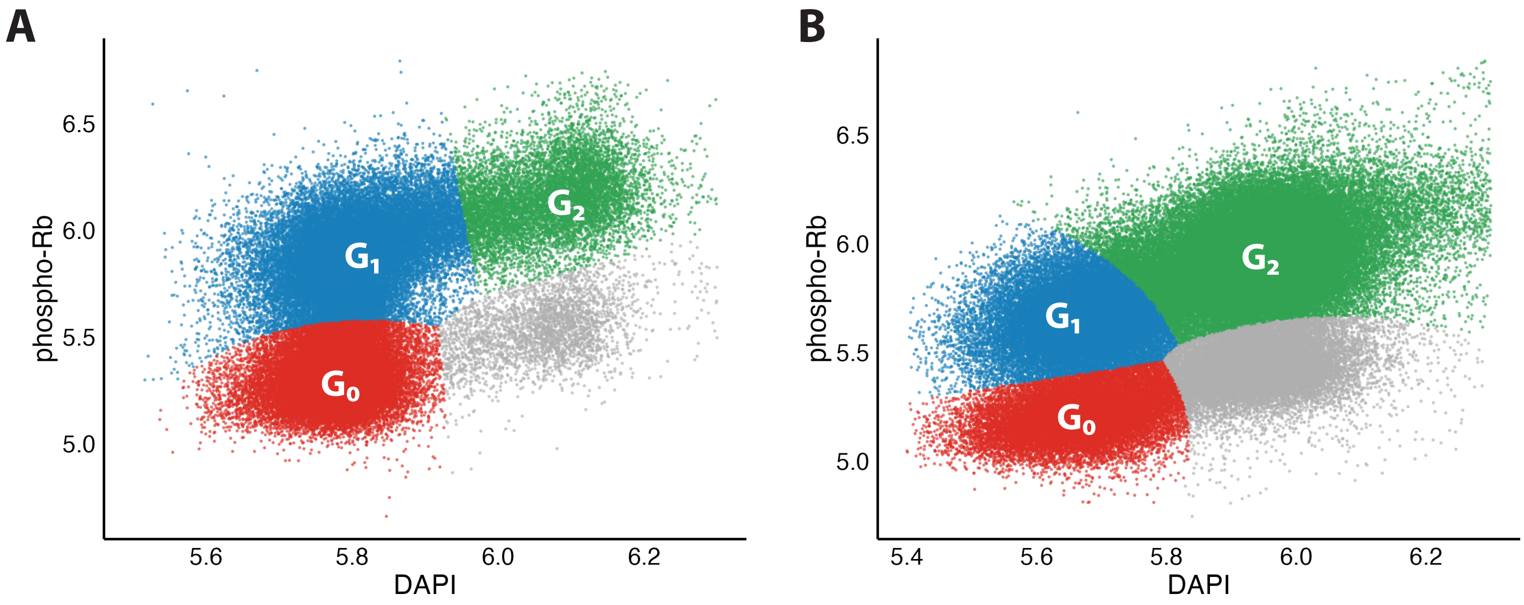
A
phospho-Rb
6.5
6.0
5.5
5.0
5.6
5.8
6.0
6.2
DAPI
G1
G2
G0
B
phospho-Rb
6.5
6.0
5.5
5.0
5.4
5.6
5.8
6.0
6.2
DAPI
G1
G2
G0

**<u>Supplementary Figure 5: Dexamethasone induces p57 mediated cell cycle arrest.</u>** Raw clusters from density-based modeling of (A) *Ptch1*$^{+/-}$ MB cells (IPM131) and (B) *Ptch1*$^{+/-}$;*Trp53*$^{-/-}$ MB cells (PTCP53 304)
